# The alveolar gas equation: do lessons from WW2 research into high flying pilots provide insights into the adaptation to high altitude flight in birds?

**DOI:** 10.1101/2020.05.13.093740

**Authors:** Michael Seear

## Abstract

Dalton’s law of partial pressures applies equally to birds and mammals so, as gas moves from the nostrils to the smallest gas-diffusion airways, the sequential addition of water vapour and CO2, steadily reduce the partial pressure of O2 (PO2) within the gas mixture. The PO2, at the point of gas exchange, at sea level, will be about 60 mm Hg less than the original PO2 within atmospheric air. As a result, the inspired PO2 is an inaccurate starting point for any model of oxygen transport. In humans, the interactions of gases at the point of diffusion, is described and quantified by the Alveolar Gas Equation (AGE). Its development during WW2, provided valuable insights into human gas exchange and also into the responses to high altitude flight in pilots but, except for an early study of hypoxia in pigeons, the AGE is not mentioned in the avian literature. Even detailed models of oxygen transport in birds omit the effect of CO2 clearance on pulmonary oxygen transfer. This paper develops two related arguments concerning the application of the AGE to birds. The first is that avian blood gas predictions, based on the theory of multicapillary serial arterialization (MSA), are inaccurate because they do not account for the added partial pressure of diffused CO2. The second is that the primary adaptation to hypobaric hypoxia is the same for both classes and consists of defending PaO2 by reducing PaCO2 through increasing hyperventilation. Support for the first is demonstrated by comparing PaO2 predictions made using the AGE, with published values from avian studies and also against values predicted by the theory of MSA. The second is illustrated by comparing the results of high altitude studies of both birds and humans. The application of the AGE to avian respiratory physiology would improve the predictive accuracy of models of the O2 cascade and would also provide better insights into the primary adaptation to high altitude flight.

## Introduction

*We live submerged at the bottom of an ocean of air.*

*Evangelista Torricelli 1644*

### Oxygen cascades of birds and mammals

The unidirectional air flow and cross-current gas exchange in avian lungs, differs significantly from the bidirectional air flow in mammalian lungs where gas exchange occurs across the walls of terminal air sacs (West J, et al, 2007). Despite these significant anatomical differences, from a functional point of view, the two respiratory systems are surprisingly similar. Whether in mammalian alveoli or air capillaries within the walls of avian parabronchi, gases are exchanged between air and the pulmonary capillary blood, across a thin diffusion membrane (Scheid P, 1979). After this step, the two cardiovascular systems are essentially the same. In each case, absorbed oxygen is mostly bound to hemoglobin within red cells and then drained to a four-chamber heart serving separate systemic and pulmonary circulations (Fedde et al, 1989). Both systems are able to sustain high levels of oxygen delivery, which allows specialist members of both classes to perform work under extreme conditions ranging from prolonged under-water dives (Meir et al, 2009), to flight at very high altitudes (Hawkes et al, 2016). Given these similarities, this study argues that lessons learned from over 150 years of research into the physiology of high altitude in humans have relevance for the understanding of the adaptation to hypobaric hypoxia in birds – particularly in the application of the alveolar gas equation (AGE) to birds. The basic hypothesis of this paper is that the long-standing theory of avian gas exchange based on multicapillary serial arterialization (Maina J, 2017), is incorrect because it does not include the effect of CO2 at the level of gas exchange. The AGE provides more accurate predictions for PaO2 at any altitude (West et al, 1980) and shows that control of alveolar CO2 through progressive hyperventilation, is as fundamental a part of the response to altitude for birds as it is for mammals (West J, 1982).

### Development of high altitude research in humans

Human high altitude research has an interesting history (West J, 2016). Torricelli was the first to demonstrate that the atmosphere has weight and, in the process, developed an accurate mercury barometer. Pascal later showed that atmospheric weight decreases with altitude. Serious high altitude research was pioneered in the 1860s by two French physiologists, Paul Bert and Denis Jourdanet (West et al, 2013). Using a steam powered vacuum pump, their decompression chamber allowed them to study the effects of altitude above 6,000 metres on a range of animals. They applied Dalton’s law of partial pressures to atmospheric air and showed that the adverse effects of high altitude were due to hypoxia, rather than low pressure, and could be reversed by breathing oxygen. In 1875, they helped Gaston Tissandier and two companions prepare for an attempt on the high altitude record of Glaisher and Coxwell who had somehow survived a flight above 11,000 meters in 1862. They were the first to carry uncompressed oxygen in a large bladder but the supply was insufficient for the crew of three; Tissandier was the only survivor (De Fonvielle W, 1875).

Pressurised bottled oxygen, developed for anesthesia, was tested in pilots during the last months of WW1 and its use allowed Finch to get within 500 meters of the peak on the 1922 British Everest expedition (Leigh J, 1974). As a comment on the adaptability of birds, it took another 30 years before Hillary and Norgay reached the summit – an altitude where many observers have reported birds flying without difficulty (or added oxygen). The greatest advances in high altitude physiology were made at the University of Rochester during WW2, by three physiologists, Rahn, Fenn and Otis (West J, 2012). While investigating the use of mask pressure-ventilation for pilots of unpressurised aircraft, they developed a mathematical model of the contents of alveolar air and its graphical equivalent, the O2-CO2 diagram (Fenn W, et al, 1946). They showed that the principal adaptation to altitude (in humans) was the hypoxic hyperventilatory response. The resulting drop in alveolar partial pressure of CO2 limits the reduction in partial pressure of oxygen with altitude.

Their conclusions were confirmed by numerous international field studies. In the English literature, those most frequently quoted include the first of three U.S. Navy hypobaric chamber studies (Project Everest 1, 1946) (Houston C, 1946), the months-long Himalayan Scientific Expedition (Silver hut expedition, 1961) (Pugh H, et al, 1962) and the American Medical Research Expedition to Everest (AMREE, 1981) (West J, et al, 1983). The last of these managed to collect alveolar gas samples during a climb of Mt Everest and showed that above about 6,000 meters, PAO2 was maintained at a bare minimum of 35-40 mm Hg by extreme hyperventilation, with PACO2 levels as low as 7 – 8 mm Hg taken by Dr Pizzo at the summit. There are, of course, many other physiological changes caused by high altitude - not least of which are the excessive loss of water and heat secondary to hyperventilation (Scott G, et al, 2008). However, in humans, the most essential adaptation, is the maintenance of a survivable PAO2 by steadily reducing PACO2 through hyperventilation.

The alveolar gas equation (AGE) was a valuable development in human respiratory physiology but, except for an early study of hypoxia in pigeons (Lutz P, et al, 1977), it is hardly mentioned in the avian literature. Even detailed models of oxygen transport in birds omit the effect of CO2 clearance on pulmonary oxygen transfer (Scott G, et al, 2006; Butler P, 2016; Lague S L, et al, 2016). Avian oxygen equilibration is based on a theory that the PO2 of pulmonary capillary blood rises steadily during its path because it constantly interacts with new infusions of fresh inspired air flowing in the opposite direction (Maina G, 2017). If this were true, the final capillary PO2 would be close to the PO2 of humidified inspired air (PIO2, about 150 mm Hg). In practice this has never been demonstrated. When measured directly in bar headed geese, the end expiratory PO2 to PaO2 gradient was positive, not negative (Scheid P, et al, 1989). Equally, whenever measured, the PaO2 of sea level birds is always in the 90 to 95 mmHg range – (far below the theoretical maximum if PIO2 were the equilibration point).

The AGE explains these findings by simply compensating for the partial pressure of CO2 that diffuses into the parabronchial air. The final equilibration point is not PIO2 but PIO2 minus PACO2, exactly as it is in mammals. The AGE also explains the primary adaptive value of the hypobaric hyperventilatory response (HVR). Although the HVR is well recognized in birds, it is usually considered a primary response to the need for higher oxygen flux at altitude (Scott G, et al, 2007) - hypocarbia is considered an incidental side effect rather than being the principal adaptive mechanism to high altitude flight by maintaining the PaO2 at a survivable minimum.

## Methods

The value of the AGE in avian respiratory physiology is demonstrated by using it to develop a stepwise model of oxygen’s passage from the atmosphere to the point where it is bound to red cells in the pulmonary capillary blood. Predicted values are then compared to published values from high altitude avian studies.

### Barometric pressure

Where values for atmospheric pressure are necessary, they are estimated using the barometric formula (Berberan-Santos M, et al, 1997) which relates the pressure of an ideal gas with molecular mass M, at height h, Pbaro(h), to its pressure at sea level, Pbaro(0):

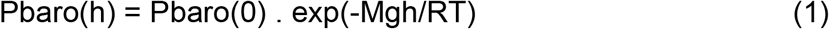

Assuming sea level pressure of 760 mmHg, a temperature of 273 deg K at sea level and constants in SI units, the equation for barometric pressure at altitude h in meters, simplifies to:

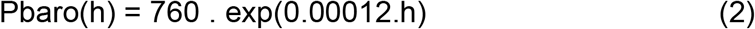

The barometric formula is considered an accurate estimate of barometric pressure in the lower stratosphere – within daily meteorological variations (Fig 1).

**Fig 1.**
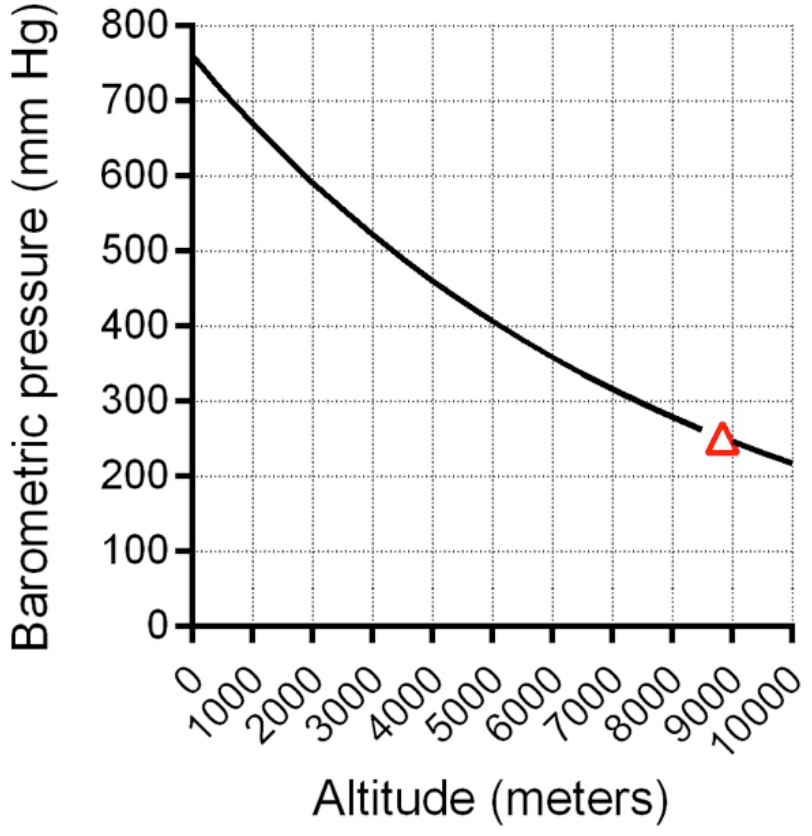
Predicted change in pressure with altitude using the barometric formula (Equn 1). M: molar mass of air (.02896 Kg/mol), g: gravitational acceleration (9.807 m/s), R: universal gas constant (8.3143 N.m/mol.K), T: standard temperature (assumed 273.15 deg K), h: altitude in meters. Open triangle: summit of Everest.

### Composition of alveolar gas

The partial pressure of oxygen steadily reduces as it moves down the diffusion gradient from atmosphere to red cell. Atmospheric air contains only two principal gases – oxygen and nitrogen. If the air is assumed to be dry, PatmO2 is simply a fraction of atmospheric pressure:

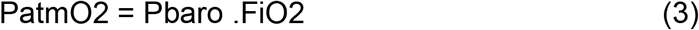

Assuming Pbaro of 760 mmHg and FiO2 of 0.21, PatmO2 at the point of inhalation is about 160 mmHg.

As air passes through the upper airway, into the trachea, a third gas (water vapour) is added. Since the gas is saturated, it dilutes the oxygen in the inspired air as follows:

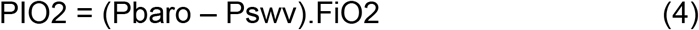

At this point, there is no difference between birds and mammals except their body temperatures. At a human’s temperature of 37 deg C, the saturated water vapour pressure (Pswv) is 47 mmHg which gives a PIO2 of 150 mmHg (Equn 4). PIO2 will be slightly lower in birds because their higher body temperature gives a higher Pswv.

As inspired humidified gas moves from the trachea into the lungs, a fourth gas is added as CO2 diffuses into the mix. In mammals, the partial pressure of alveolar oxygen at steady state, is predicted by the alveolar gas equation (Subramani S, et al, 2011). There is a small added factor which affects the final value of PAO2 by less than 1% so the simplified equation is usually used:

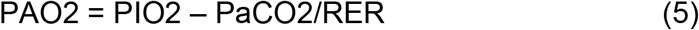

Assuming an arterial CO2 partial pressure (PaCO2) of 40 mmHg and a Respiratory Exchange Ratio (RER) of 0.8, the PAO2 in a healthy human at sea level is about 100 mmHg. In mammals, there is an oxygen gradient from alveolus to arterial blood due to small shunts and minor V/Q mismatch. In healthy non-smokers, PAO2-PaO2 gradient is 5 to 10 mmHg. In birds, the situation is less clear. The gas flux across bird lungs is certainly more efficient than it is in mammals but birds cannot break Dalton’s law of partial pressures. Pbaro at a given altitude is fixed. If CO2 is added to the mix within the parabronchi, then the partial pressure of O2 must fall. The resulting blood gases are better predicted using the AGE than they are by theories based on multicapillary serial arterialization (MSA).

### Changes of alveolar gases with altitude

As gas moves from nostrils to the smallest gas-diffusion airways, the sequential addition of water vapour and CO2, reduces the PO2 by about 60 mm Hg. As a result, the PIO2 is an inaccurate starting point for models of oxygen transport at any altitude. The PO2 in all three inspiratory compartments falls predictably with altitude (Fig 2). By the summit of Everest (8840 m), Pbaro falls to 252 mmHg (equn 2). Similarly, PatmO2 will be 53 mmHg (equn 3) and PIO2 will be 43 mmHg (equn 4). If PCO2 is not included then this gives a false impression of the PO2 available at that altitude. Assuming a climber with an RER of 0.8 and a PaCO2 fixed at 40 mmHg, the AGE (equn 5) shows that PAO2 will fall to zero long before the summit is reached (Fig 2). Without added oxygen, it is simply impossible to survive at high altitude without sustained hyperventilation.

**Fig 2.**
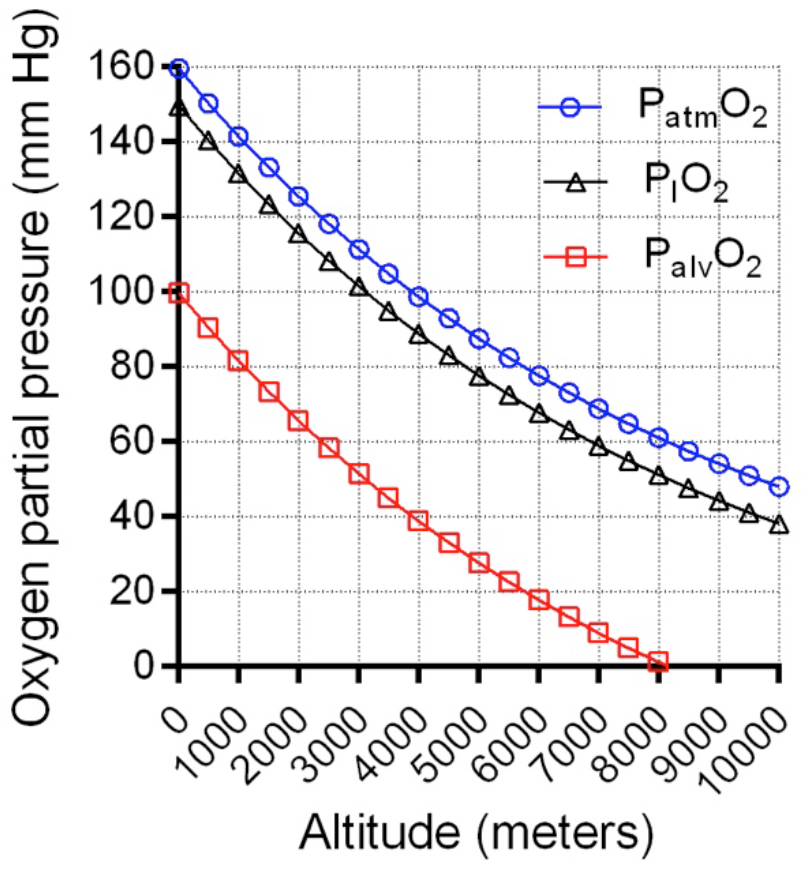
Predicted effects of altitude on the partial pressures of oxygen in three respiratory compartments - dry atmospheric air: PatmO2 (equn 3), humidified inhaled air: PIO2 (equn 4), and alveolar gas: (equn 5), assuming PaCO2 of 40 mm Hg and RER of 0.8.

### Effect of the hyperventilatory response

In order to maintain a survivable PaO2, the PaCO2 must fall steadily with altitude. As noted, measurements taken during the 1981 American Medical Research Expedition, showed that the PaCO2 in a climber on the summit, was below 10 mm Hg (West J, et al, 1983). Substituting this into the AGE gives the climber a bare minimum PAO2 around 35 mmHg. The ability to reach the summit without bottled oxygen is so marginal that a reduction in Pbaro of only 5 mmHg would probably make the feat impossible. Figure 3 shows how lowering PaCO2 by hypobaric hyperventilation is able to modify the reduction in PAO2 with altitude. The PaO2 of a climber is superimposed showing how increasingly extreme hyperventilation maintains a survivable PaO2, particularly above about 6,000 meters. Birds appear to tolerate combinations of hypoxia and hypocarbic alkalosis that would cause rapid unconsciousness in humans due to cerebral vasoconstriction (Butler P J, 2010). With slow ascent over several days, humans can increase renal excretion of bicarbonate, which returns arterial pH to normal levels allowing then to tolerate low PaCO2.

**Fig 3.**
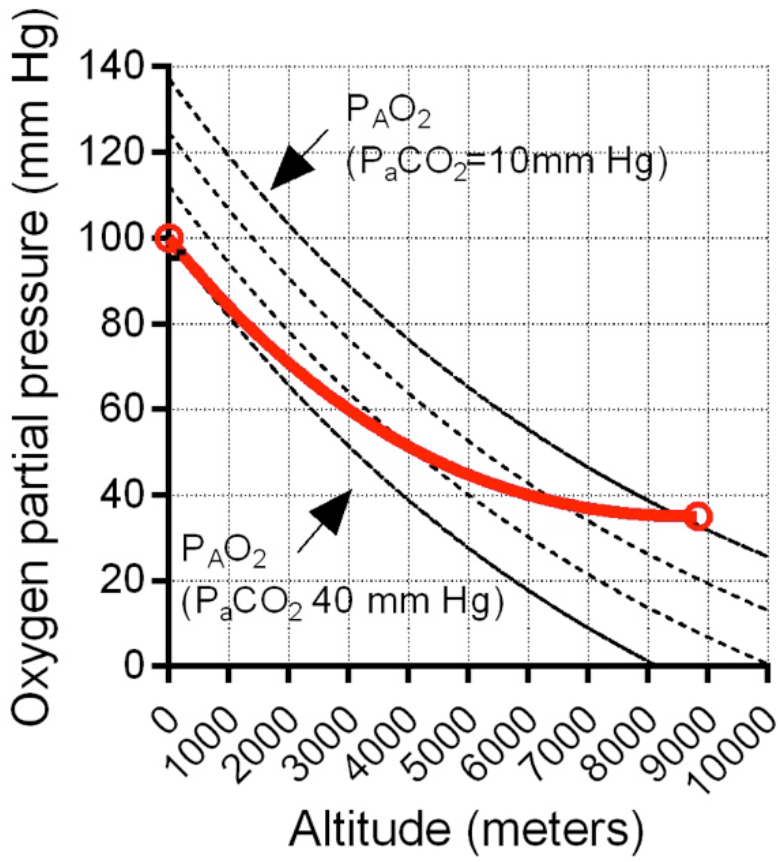
Dotted lines - predicted changes of PAO2 with altitude, calculated for four values of PaCO2 (AGE equn 5). RER assumed to be 0.8 in all cases. The solid line shows the progressive decrease in PAO2 of a climber going from sea level (PAO2 100 mmHg), to the summit of Everest (PAO2 35 mmHg). The climber’s path crosses PAO2 lines as adaptive hyperventilation reduces PaCO2 to below 10 mm Hg at the summit of Everest.

### Hemoglobin oxygen saturation and blood oxygen content

Less than 2% of the oxygen carried in arterial blood (CaO2) is dissolved in plasma – the bulk is carried on hemoglobin (Hgb). In mammals, arterial oxygen content (CaO2) is described by the expression:

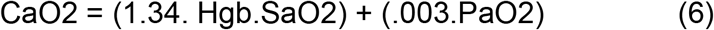

Oxygen saturation (SaO2) is expressed as a fraction and results are given as ml O2 per 100 ml blood. Assuming Hgb of 15, SaO2 of 1.0 and PaO2 of 100 mm Hg, each 100 ml of arterial blood contains 20.1 ml O2 attached to Hgb and 0.3 ml of O2 in solution (equn 6). This equation has been used in avian studies and shown to produce results within 5% of measured values in penguins (Meir J U, 2009). The main variables determining blood oxygen content are clearly Hgb concentration and the affinity of Hgb for oxygen saturation. While Hgb may slowly increase in response to chronic hypoxia (Storz J, et al, 2008), rapid adaptation to oxygen needs is provided by the constantly changing affinity of Hgb for oxygen as it moves from lung to peripheral tissues (Meir J U, 2013). The hypocarbic, alkalotic and relatively cool environment at the pulmonary diffusion membrane produces a leftward shift in the oxy-hemoglobin dissociation curve (Fig 4, top) which increases Hgb affinity for O2. The warmer, more acidotic and hypercarbic environment within the vascular bed of active muscles, decreases affinity (right shift, Fig 4) and facilitates unloading of oxygen where it is needed. The shifts are only a few mm Hg in either direction but, at the steepest part of the curve, this can have a significant effect on Hgb saturation and blood oxygen content when PO2 is low (Fig 4, bottom).

**Fig 4.**
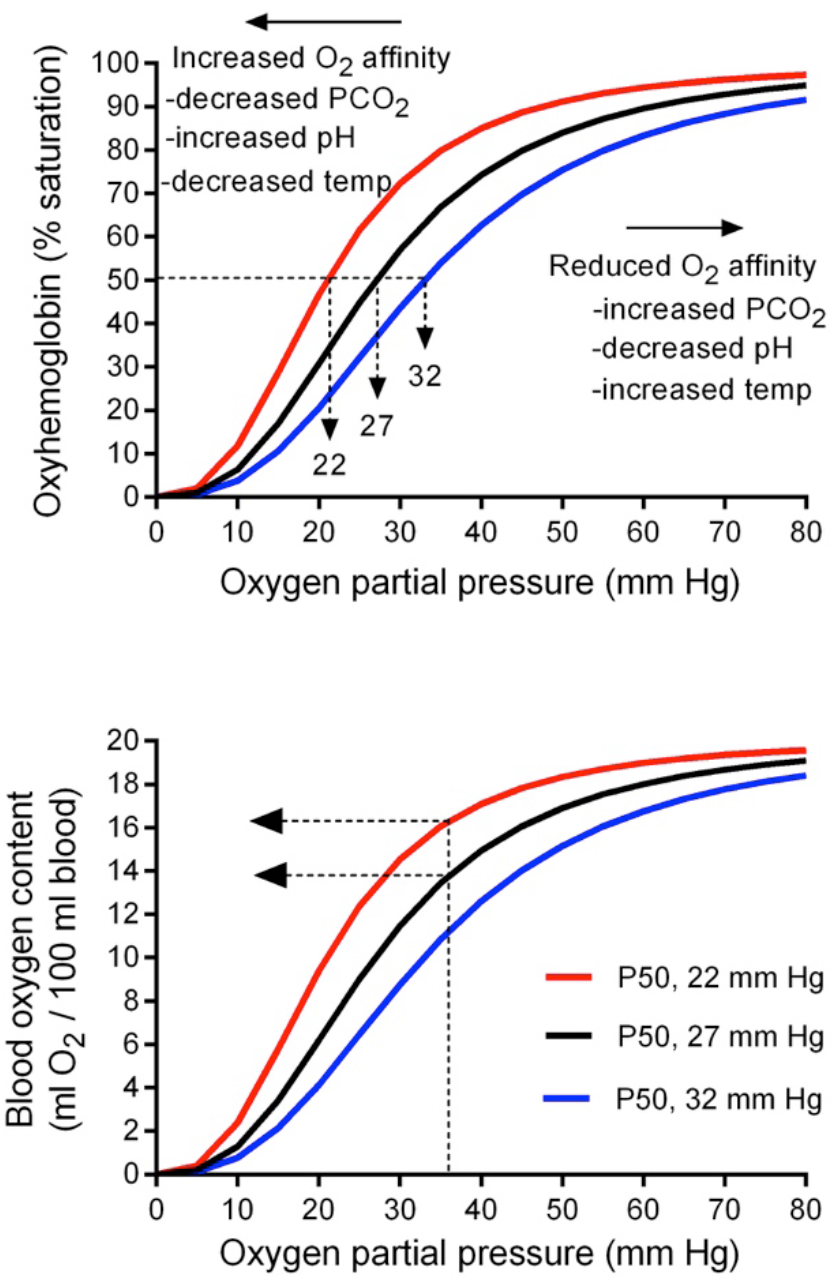
top: dissociation curves for an animal with P50 of 27 mm Hg and Hill coefficient of 2.8, demonstrating how O2 affinity varies with changes in temp, pH and PCO2. Fig 4, bottom: the effect of a 5 mm Hg left shift on O2 content for a climber at the summit of Everest with PAO2 of 35 mm Hg and CaO2 about 14 ml/100 ml (equn 6). CaO2 increases over 15% allowing bare minimum work at a survivable SaO2 of 80%.

There are many equations available to model the oxy-hemoglobin dissociation curve. The following has been well tested (Dash R, et al, 2016):

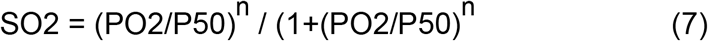

P50 is the PO2 that results in 50% Hgb saturation and n is the Hill coefficient, which is a measure of the steepness of the central part of the dissociation curve. Avian models usually use a value of 2.8 (Scott G, et al, 2006), which is the same for humans and birds. Several studies have shown that the P50 is generally higher in birds than mammals but high altitude specialists, such as bar-headed geese, have developed a left shift towards a P50 of 27 mm Hg, the same as humans (Lutz P L, 1980).

## Results

Two related theoretical arguments have been developed in ‘Methods’. The first is that avian blood gas predictions based on the theory of multicapillary serial arterialization (MSA) are inaccurate because they do not account for the added partial pressure of diffused CO2. The second is that the primary response to hypobaric hypoxia is the same for both classes and consists of defending PaO2 by reducing PaCO2 through increasing hyperventilation. Support for the first is demonstrated by comparing PaO2 predictions made using the AGE, with published values from avian studies and also against values predicted by the theory of MSA. The second is illustrated by comparing the results of high altitude studies of both birds and humans.

### • Prediction comparisons, accuracy and potential errors

It should be remembered that high altitude research is very difficult. Whether performed in a hypobaric chamber or in a remote field location, measurement accuracy is a challenge. Problems include measurement errors of portable gas analysers (Harter T, et al, 2016) and failure of the animal to reach steady state under extreme conditions. Comparisons between the results of different studies and subsequent conclusions, must be interpreted with caution.

Table 1 displays predictions of SaO2 and CaO2 for bar-headed geese (BHG) at 7,000 meters using the equations developed in Methods above. Values in the left column are calculated using the AGE, those in the right are based on the theory of MSA which does not adjust for PCO2 within the small airways. This over-estimates PAO2, SaO2 and calculated blood oxygen carrying capacity (right column, table 1). The assumptions made in the equations are based on published values for BHG (Black C, 1980: Hawkes L, 2014; Faraci F, 1984). In reality, under extreme conditions, these variables are probably in constant flux so the assumption of steady state is not necessarily accurate. Even small changes in variables such as RER, PaCO2 or shift of the dissociation curve, can cause significant changes in CaO2.

**Table 1.**
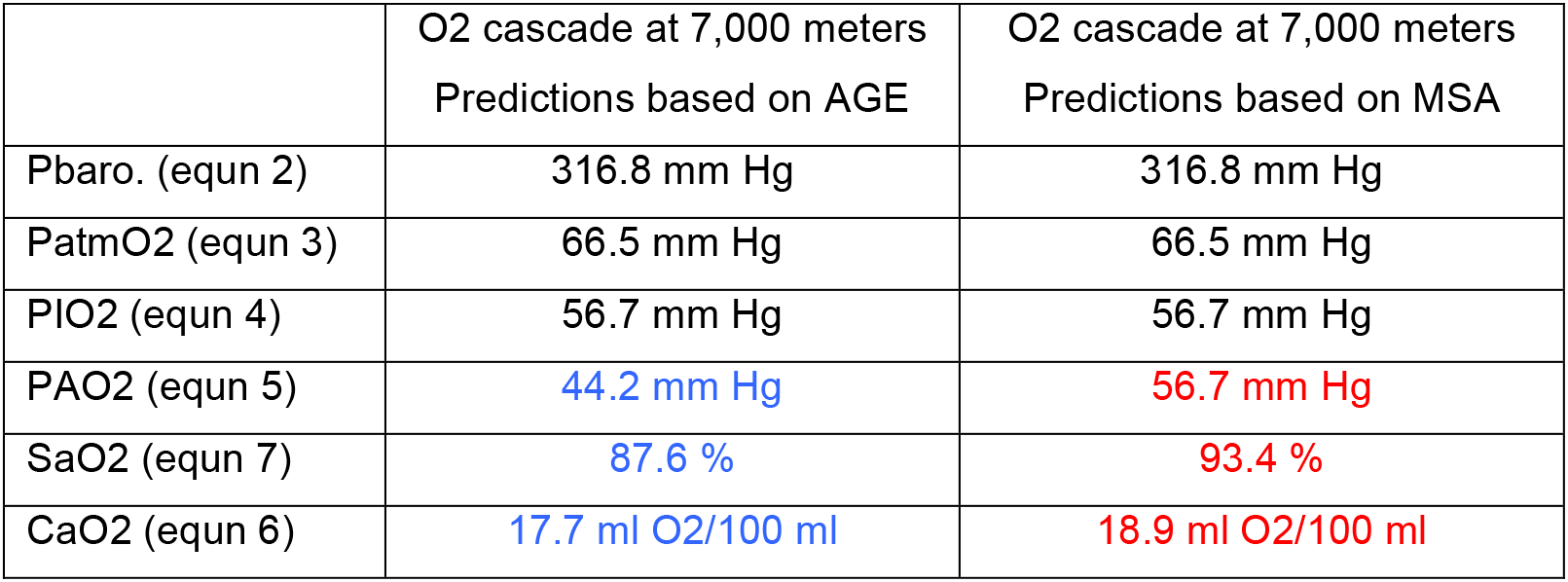
Predicted values for steps in the oxygen cascade from inspired atmospheric air to hemoglobin saturation, at 7,000 m altitude, using equations 2 to 7. Values in left column calculated using AGE (Alveolar Gas Equation), those in right column calculated using MSA (Multicapillary Serial Arterialisation). Assumptions for both are based on published high altitude studies of bar-headed geese: body temp 41 deg C, Pswv 58 mm Hg, Hgb 15 gm/100 ml, oxy-hemoglobin P50 of 27 mm Hg, 5 mm Hg left shift of dissociation curve, PaCO2 of 10 mm Hg at 7,000 meters.

|  | O2 cascade at 7,000 meters<br>Predictions based on AGE | O2 cascade at 7,000 meters<br>Predictions based on MSA |
| --- | --- | --- |
| Pbaro. (equn 2) | 316.8 mm Hg | 316.8 mm Hg |
| PatmO2 (equn 3) | 66.5 mm Hg | 66.5 mm Hg |
| PIO2 (equn 4) | 56.7 mm Hg | 56.7 mm Hg |
| PAO2 (equn 5) | 44.2 mm Hg | 56.7 mm Hg |
| SaO2 (equn 7) | 87.6 % | 93.4 % |
| CaO2 (equn 6) | 17.7 ml O2/100 ml | 18.9 ml O2/100 ml |

A few studies – both in birds and humans, give details of arterial blood gas values at stated altitudes. This allows PaO2 to be calculated using the AGE (equn 5). Those values are compared to measured values of PaO2 in Fig 5. The result can only be viewed as an approximation since the plot relies on pooled average data from six studies. Despite this, the correlation between theory and practice is excellent (R>0.9), and supports the basic assumptions of this paper – birds and humans are both governed by the law of partial pressure and the best predictor of alveolar gases in both classes is the AGE. If PIO2 were the equilibration point for parabronchial gases then PAO2 would be close to 150 mm Hg but the highest published value for birds at sea level is never above 100 mm Hg (Fig 5).

**Fig 5.**
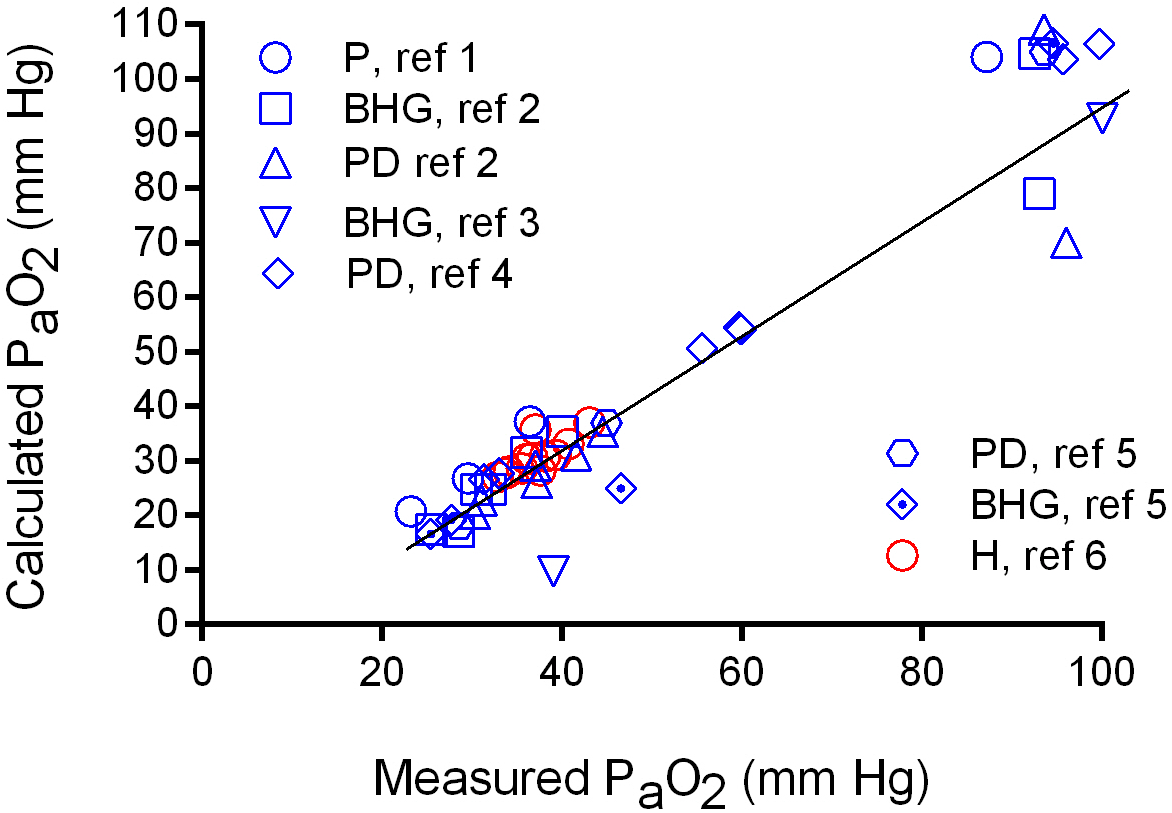
Pooled values for PaO2 from six publications, plotted against calculated values using the AGE (equn 5). Regression line calculated using method of least squares, correlation coefficient (R) >0.9. P: pigeon, BHG: bar headed goose, PD: pekin duck, H: human. Ref 1: (Lutz P, et al, 1977), ref 2: (Black C, et al, 1980), ref 3: (Hawkes L, et al, 2014), ref 4: (Shams H, et al, 1990), ref 5: (Faraci F, et al, 1984), ref 6: (West J, et al, 1983). P_A_O_2_ to P_a_O_2_ gradient assumed to be 5 mm Hg. Note that no measured value of PaO2 is higher than 100 mm HG.

### Comparison of high altitude responses in birds and humans

The relationship between barometric pressure and the partial pressures of respiratory gases is tightly controlled by the law of partial pressures and expressed mathematically by the alveolar gas equation. Values of PaO2 and PaCO2, in a mixture of non-reactive gases, are dependent on each other – if one rises, the other must fall. Results are best visualized by plotting simultaneous values (measured at steady state), to produce an O2 – CO2 diagram. This is a convenient way to compare the hyperventilatory adaptations to high altitude in birds and mammals.

With the same precautions concerning potential errors, the data from the six studies used in Fig 5, can also be used to construct an O2-CO2 diagram showing how these two gases change during ascent to extreme altitude (Fig 6). Starting at sea level, PaO2 initially falls steadily but, above about 6,000m, increasing hyperventilation limits the rate of decline by steadily reducing PaCO2 values to very low levels. The solid line in Fig 6 is based on human data collected during the 1981 AMREE study and shows that the altitude response in humans and birds is identical. Both defend PaO2 by reducing PaCO2 to very low levels through increasingly severe hyperventilation.

**Fig 6.**
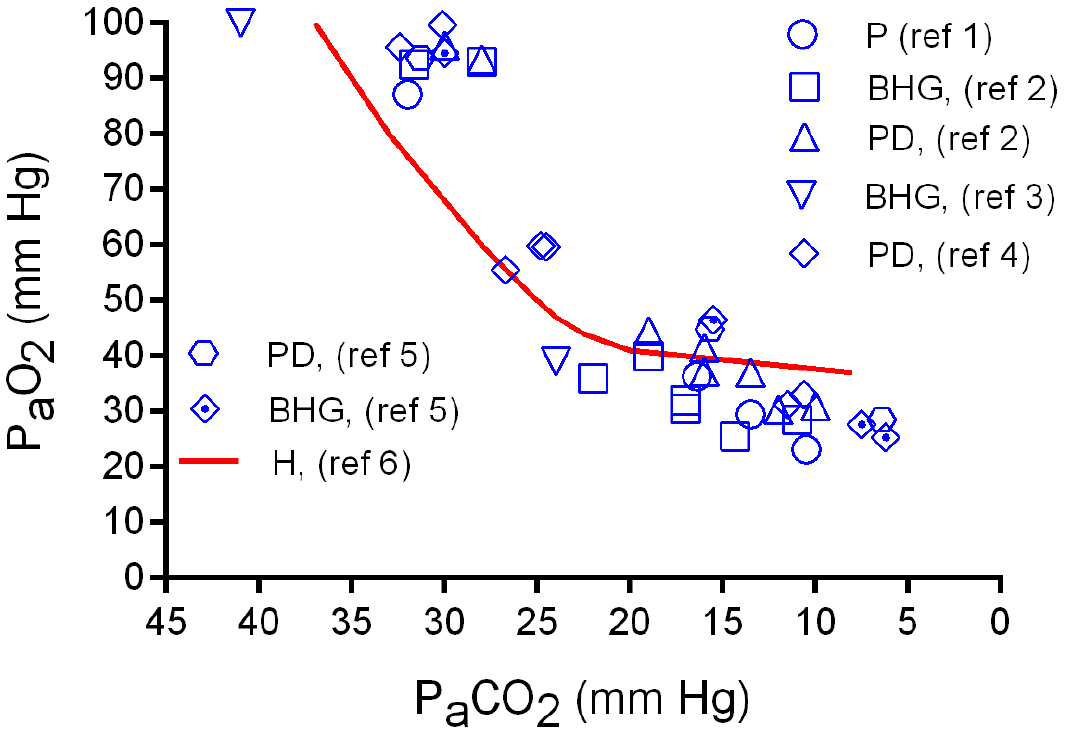
Pooled values for arterial blood gases from six high altitude studies. PaO2 plotted against simultaneous values for PaCO2. P: pigeon, BHG: bar headed goose, PD: pekin duck, H: human. Ref 1: (Lutz P, et al, 1977), ref 2: (Black C, et al, 1980), ref 3: (Hawkes L, et al, 2014), ref 4: (Shams H, et al, 1990), ref 5: (Faraci F, et al, 1984), ref 6: (West J, et al, 1983). Blue symbols are avian studies. Solid red line represents human altitude adaptation (ref 6).

## Discussion

This paper has developed the proposal that the theory of avian gas exchange, based on serial stepwise oxygenation within the pulmonary capillaries, is inaccurate because it does not adjust for the partial pressure of added CO2 at the point of gas diffusion. Birds and mammals have very different lungs but both must follow Dalton’s law of partial pressures. Using published data, comparison between values of PaO2 calculated with the AGE and measured values, showed excellent correlation (Fig 5) confirming that the AGE applies equally to birds and mammals and provides a more accurate starting point for models of the oxygen cascade in both classes. More importantly, the AGE provides insight into the importance of the HVR as the primary adaptation to high altitude in birds. At constant Pbaro, values for PACO2 and PAO2 are tightly bound – if one is reduced, the other rises. This means that the reduction in PAO2 with altitude can be minimized, within limits, by steadily reducing PACO2 by hyperventilation. Using published high altitude studies of birds and humans, figure 6 illustrates this adaptation and shows that the response applies equally to birds and mammals.

Research into the flow patterns and interactions of gas and blood within avian lungs is difficult because both are complex and still incompletely defined. For example, gas flow through the paleopulmonic parabronchi is unidirectional, but it is believed to be tidal within neo-pulmonic parabronchi (Weidner W, 2011). Similarly, pulmonary capillary blood flows perpendicularly to gas flow within the parabronchi but it runs directly counter to gas flow within the air capillaries (Maina J, 2017). Whether the resulting gas exchange is best described as cross-current, counter-current, or a mixture of both, was debated at length in earlier literature (Scheid P, et al, 1972). The theory that emerged, termed multicapillary serial arterialization (MSA), claims that capillary blood moves counter to a continuous flow of inspired air so it is steadily exposed to fresher gas that hasn’t yet given up its O2. If this were true, the final equilibration point for pulmonary capillary blood would be the PO2 of inspired humidified air, which is roughly 150 m Hg. This is easily disproved – the PaO2 in any study of birds or humans at sea level is never 150 mm Hg but is in the range of 95-100 mm Hg; the data displayed in figures 5 and 6 provide typical examples.

The difference between PaO2 predictions using the AGE and the theory of MSA is due to the PACO2 which equals PaCO2/RER (equn 5). The value is about 40/0.8 mm Hg at sea level. While this causes an appreciable error in PaO2, the shape of the dissociation curve limits the effect of this error when calculating CaO2. Hemoglobin is fully saturated at a PaO2 of 100 mm Hg. A 50 mm Hg increase in PaO2 above this level would all be dissolved in plasma and only increase CaO2 by an extra 0.3 ml/100 ml blood (equn 6). At altitude, the situation is different. As PaCO2 decreases with altitude, the difference between predicted values for PaO2 also becomes smaller. By 7,000 m, it is only 10 to 15 mm Hg (Table 1). The theory of MSA still overestimates CaO2 but the difference is small (Table 1). The AGE is a more accurate representation of the physiology of gas exchange so is a better foundation for mathematical models of any aspect of the oxygen cascade in mammals or birds.

The value of adapting the AGE to birds is only partly due to an improved ability to model blood oxygen content and delivery. Its greater advantage is in providing better insights into the acute adaptations to high altitude flight. Based on this paper’s findings, the main responses to the challenges of high altitude are listed in table 2. It is worth starting with the importance of route finding. While BHG have been spotted flying above Everest, the low PO2 and air density at those heights make it incredibly challenging. Not surprisingly, tracker studies have shown that birds choose routes that allow them to stay well below 6,000 m most of the time (Hawkes L A, et al, 2013). The main physiological response to hypobaric hypoxia is the defence of PaO2 by progressively reducing PaCO2. This is not mentioned in the avian literature. The HVR is recognized but is said to be due to the need for greater O2 flows at altitude. While increased O2 flux is necessary, respiratory drive is also governed by the need to defend a survivable PaO2. The AGE clearly shows that without hyperventilation, the PaO2 would steadily fall to zero, long before the top of Everest (Fig 2). Hypocarbia is not just an incidental side effect of hyperventilation but is the primary adaptation that makes high altitude flight possible.

**Table 2.**
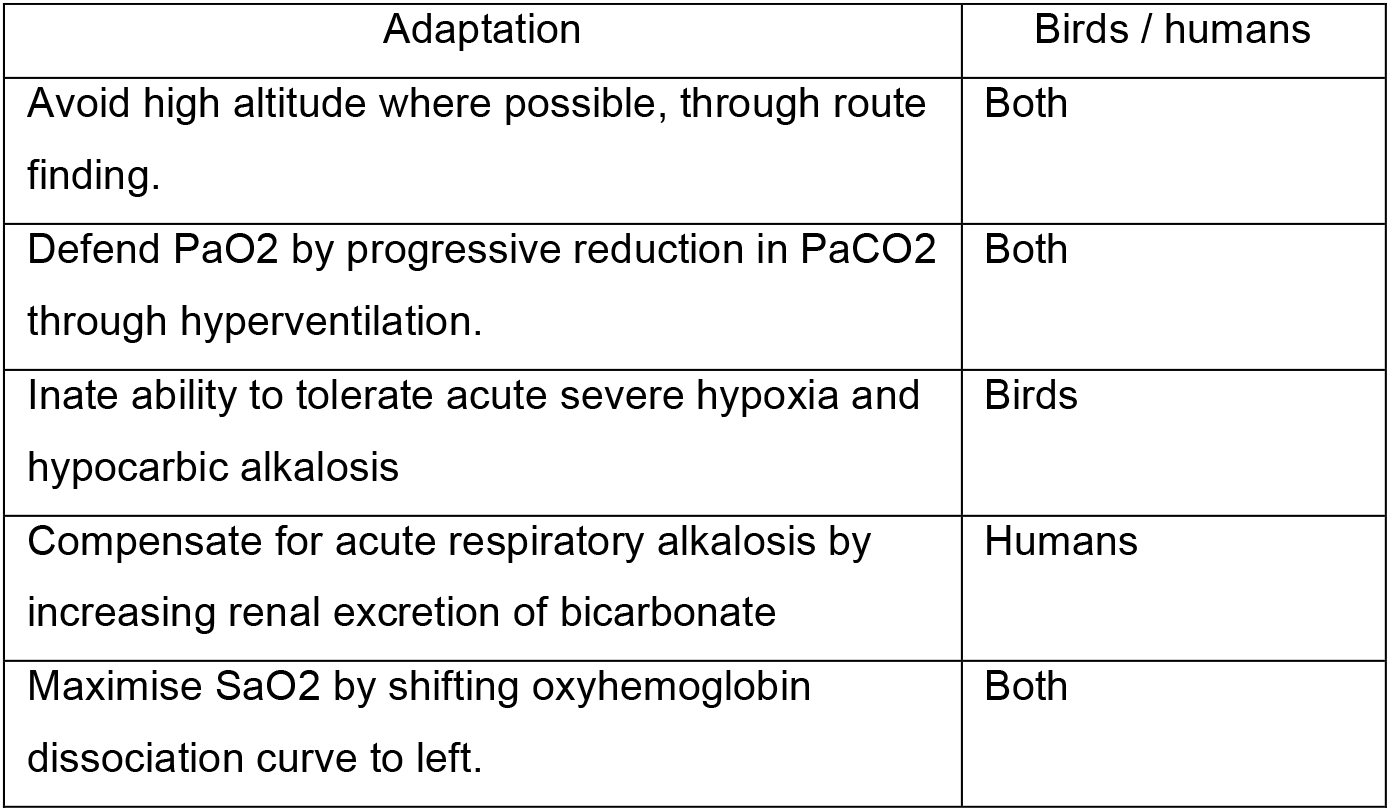
Principal adaptations to the challenges of high altitude.

| Adaptation | Birds / humans |
| --- | --- |
| Avoid high altitude where possible, through route finding. | Both |
| Defend PaO<sub>2</sub> by progressive reduction in PaCO<sub>2</sub> through hyperventilation. | Both |
| Inate ability to tolerate acute severe hypoxia and hypocarbic alkalosis | Birds |
| Compensate for acute respiratory alkalosis by increasing renal excretion of bicarbonate | Humans |
| Maximise SaO<sub>2</sub> by shifting oxyhemoglobin dissociation curve to left. | Both |

Since controlled hypocarbia is not recognized as an altitude adaptation in birds, many alternative explanations have been suggested (Parr N, et al, 2019). In terms of relative importance, it should be remembered that those various responses all share a common factor – the bird has to be alive and that requires a survivable PaO2, which in turn, requires hyperventilation. The associated acute hypocarbic alkalosis is tolerated by birds but not humans. They require a separate renal compensation for the alkalosis before hyperventilation can be tolerated. The final acute adaptation is the transfer of a low PaO2 into a level of saturation that can support work. This is achieved by a left shift of the dissociation curve. The oxygen cascades and adaptations to altitude in humans and birds are more similar than has previously been accepted. This paper has hopefully shown that the O2 cascade can be represented by a series of physiological equations that are applicable to both classes of animal, including the AGE. There is a need for research targeted at defining more accurate values for the numerous constants included within those equations.

## Abbreviations

P: partial pressure
O2: oxygen
CO2: carbon dioxide
PatmO2/CO2: atmospheric partial pressure of O2 or CO2
PIO2/CO2: humidified inspired partial pressure of O2 or CO2
PAO2/CO2: alveolar partial pressure of O2 or CO2
PaO2/CO2: arterial partial pressure of O2 or CO2
Pswv: saturated water vapour pressure
Pbaro: barometric pressure
FiO2: fraction of inspired O2
RER: respiratory exchange ratio
MSA: multicapillary serial oxygenation
Hgb: hemoglobin
SaO2: saturation of arterial O2
n: Hill coefficient
CaO2: arterial O2 contentP50
P50: PaO2 producing 50% Hgb saturation
HVR: hypoxic ventilatory response

